# Effects of METTL3-METTL14 on primary microRNA processing by Drosha-DGCR8

**DOI:** 10.1101/2024.10.15.618347

**Authors:** Brijesh Khadgi, Yunsun Nam

## Abstract

MicroRNAs modulate most protein-coding genes, and many are regulated during maturation. Chemical modifications of primary transcripts containing microRNAs have been implicated in altering Microprocessor processing efficiency, a key initiating endonucleolytic step performed by Drosha and DGCR8. METTL3-METTL14 produces N^6^-methyladenosine which is the most common methylation for mRNAs. Genetic experiments suggested that METTL3-METTL14 promotes primary microRNA processing by Microprocessor, but the molecular mechanism still needs to be elucidated. We tested the hypothesis that METTL3-METTL14 or m^6^A may directly impact Drosha or DGCR8 function during primary microRNA processing. After reconstituting the methyltransferase and processing activities, we show that the presence of METTL3-METTL14 complexes does not affect the processing efficiency of Drosha-DGCR8. We also established a method to prepare m^6^A-modified primary microRNAs and used them to show that the processing of the transcripts with m^6^A is similar to those without any modification. Recombinant METTL3-METTL14 and DGCR8 do not form stable complexes, challenging the previous model that depends on enhanced DGCR8 recruitment. Therefore, METTL3-METTL14 or m^6^A modification does not generally promote Microprocessor-mediated microRNA processing, although they may impact certain cases.

## Main Text

MicroRNAs are small RNAs that can regulate specific target gene expression and are critical for proper development and maintenance of multicellular organisms (1). MicroRNAs are transcribed as part of longer transcripts (primary microRNAs or pri-miRs). To activate the ability to switch off downstream genes, microRNAs require excision steps carried out by the dedicated processing enzymes—Drosha-DGCR8 complex (termed Microprocessor) followed by Dicer (2-6). The first cleavage performed by Microprocessor is critical in determining the active microRNA function, and mature microRNA levels do not correlate well with the abundance of the primary transcripts (7,8). Many key signaling pathways control gene expression patterns by regulating microRNAs during the Microprocessor-dependent processing step (9).

MicroRNA processing may also be regulated via chemical modifications of the primary transcript. Coding and noncoding RNAs are decorated with various modifications post-transcription, which can regulate almost every aspect of the RNA life cycle (10,11). Two modifications have been highlighted for their impact on Microprocessor-dependent processing of microRNAs. N^6^-methyladenosine (m^6^A) modification by METTL3-METTL14 is the most abundant internal modification found in mRNAs (12,13). Primary microRNAs can also contain m^6^A modifications (14), and numerous studies have noted that increased expression of the METTL3-METTL14 methyltransferase complex and higher m^6^A levels correlate with more efficient microRNA maturation, resulting in increased mature microRNA levels in cancer cells of various types such as in the bladder, pancreas, intestines, liver, lung, and breast (15-23). Another modification, internal N^7^-methylguanosine (m^7^G), can be added by the METTL1-WDR4 complex. m^7^G has been implicated in promoting the maturation of key pri-miRs such as those of the let-7 family of tumor-suppressive microRNAs (24). Modifications have been linked to better recruitment of Microprocessor via DGCR8 (15,22,25) or remodeling of obstructive G-quadruplex structures in the pri-miRs to promote processing (24). However, previous studies were performed in the presence of other cellular factors, making it difficult to determine the direct impact of the methyltransferase enzymes and chemical modifications on the primary transcripts and their processing.

Here we show that primary microRNAs can be readily modified with m6A but not m7G, using reconstituted METTL3-METTL14 and METTL1-WDR4 enzymes. The heterodimeric METTL3-METTL14 complex robustly methylates many pri-miRs, but does not positively or negatively impact primary microRNA processing by recombinant Drosha-DGCR8. We established a new method to purify transcribed primary microRNAs with m6A modification by using an m6A-specific reader protein, YTHDC1. Using the well-defined RNA substrates, we established that primary microRNAs are processed similarly by Drosha-DGCR8, in the presence and absence of m6A. The processing rates also do not change in the presence of saturating levels of YTHDC1, and METTL-METTL14 do not interact with purified DGCR8. Therefore, we conclude that METTL3-METTL14 and the m6A modification do not directly impact Microprocessor activity.

## Results

To investigate how RNA methyltransferases affect microRNA processing, we set out to reconstitute the methylation and processing enzymes. We purified METTL3-METTL14 and METTL1-WDR4 as heterodimer complexes as METTL14 and WDR4 are needed for methyltransferase activity for METTL3 and METTL1, respectively (26,27). We used pri-let-7e as the substrate RNA as it has been shown to be methylated by both enzymes (14,24). METTL3-METTL14 showed robust methylation activity with the pri-miR, but incubating with METTL1-WDR4 did not show any detectable methylation (Figure 1A, B). In contrast, METTL1-WDR4 showed high methylation activity with tRNA^Trp^, indicating catalytic competence. Structures of METTL1-WDR4 in complex with tRNA suggested that the methyltransferase recognizes the tRNA fold to target G46 for methylation (27). Most pri-miRs have structures distinct from tRNAs. Furthermore, since the methylation activity of METTL1-WDR4 with pri-miR substrates was too low to be detectable, most pri-miRs are not likely to be modified by METTL1. Therefore, due to its more robust methylation activity on pri-miRs, we focused our study on investigating the effects of METTL3-METTL14 on microRNA maturation.

**Figure 1.**
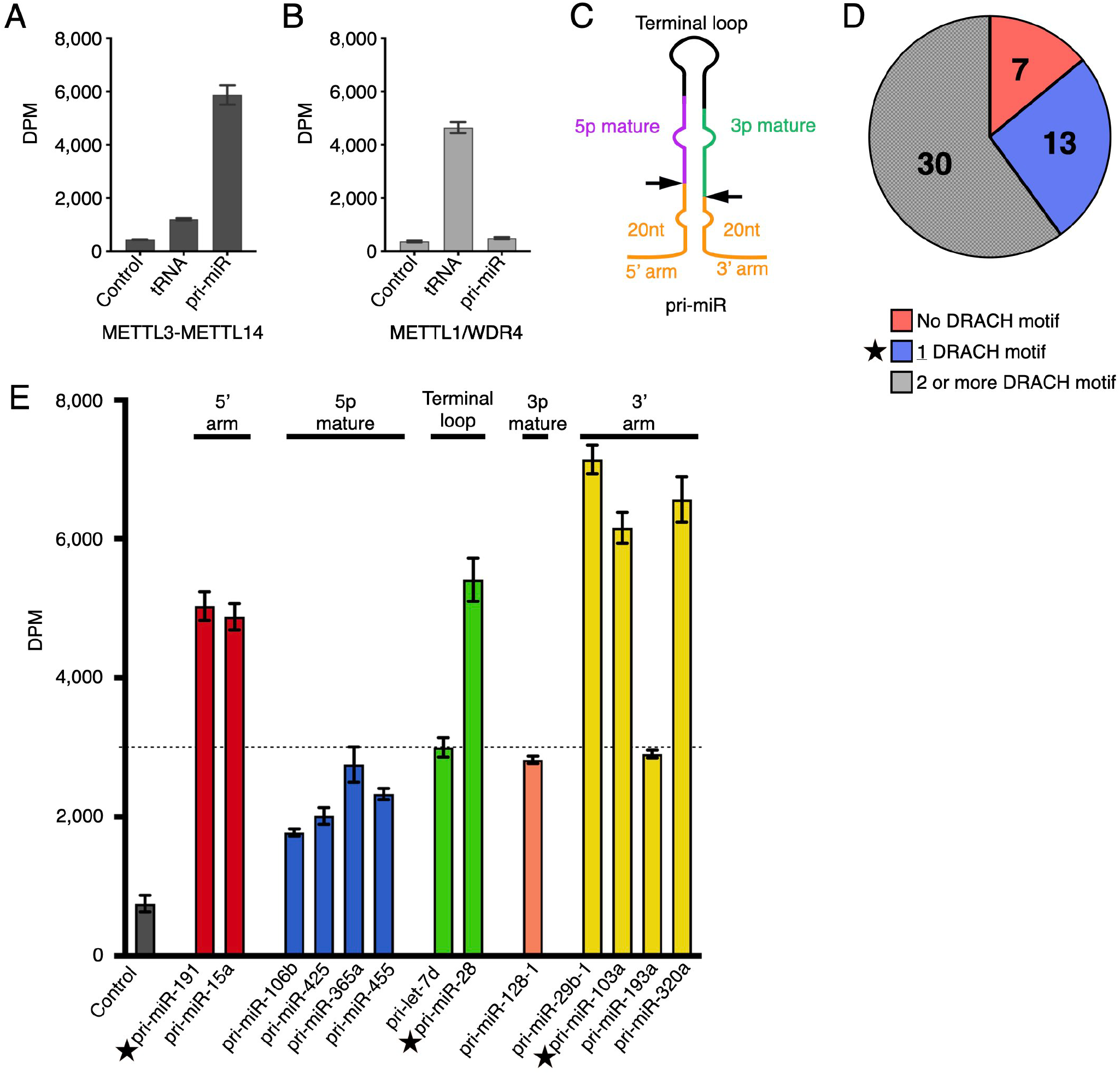
METTL3-METTL 14 robustly methylates pri-miRs. **(A-B)** In vitro methylation assays show that METTL3-METTL 14 methylates pri-let-7e (pri-miR) but not tRNATrP **(A)**, whereas METTL 1-WDR4 methylates tRNATrP but not pri-let-7e **(B)**. Tritiated (^**3**^H) methyl groups transferred from SAM to RNAs were quantified by scintillation counting measured by disintegrations per minute (DPM). **(C)** Schematic representation of a minimal pri-miR substrate for Microprocessor. The hairpin structure contains a mature microRNA on either strand (purple or green). The 5’ and 3’ arms (orange) extend beyond the Microprocessor cut sites (arrows), and 20 nucleotides on each side are sufficient for a Microprocessor substrate. **(D)** 50 pri-miRs show sensitivity to METTL3 knock-down (fold change < 0.5 and p value of< 0.05). The pie graph shows the pri-miRs with different numbers of DRACH motifs, limited to arm lengths of 20 nt on each side. **(E)** In vitro methylation assays show varying methylation propensities for pri-miRs with only one consensus methylation motif from **(D)**. The bars are grouped by the location of the DRACH motif (specified on top and by color). Three readily methylated pri-miRs (starred) were chosen for further analysis.

Many mature microRNA levels decrease upon knocking down METTL3 (14). To identify the key microRNAs to investigate in depth, we focused on the group of microRNAs of which mature levels decreased by more than 2-fold (p < 0.05) upon knocking down METTL3; this group contains 50 pri-miRs (Supplementary Table S1). METTL3-METTL14 is known to preferably methylate a specific sequence containing DRACH (28,29). We looked for the consensus motif among the 50 pri-miRs. For each pri-miR, Microprocessor interacts with the flanking arms beyond the Drosha cut site, and 13 nucleotides on each side of the cut site have been shown to be minimally required for full activity (30). Thus, we limited the consensus motif search to the pri-miR with the flanking arms of 20 nucleotides in length (Figure 1C).

Among the pri-miRs that contain at least one DRACH motif in the Microprocessor footprint (20-nt away from the cut sites), we selected those that contain only a single DRACH motif, because multiple m^6^A sites would likely confound any position-specific regulatory effects (Figure 1D). We generated purified pri-miRs for all 13 of the microRNAs that contain exactly one DRACH motif. These pri-miRs show varying propensities for m^6^A modification, likely due to structural constraints (Figure 1E) (26). Our results indicate that METTL3-METTL14 most efficiently methylates DRACH sequences that are more likely to be in duplexes, such as the flanking arm regions or the terminal loop. Furthermore, certain pri-miRs are more prone to erroneous processing, making the processing efficiency difficult to quantify. Thus, we chose three pri-miRs with higher methylation propensity and more precise processed products, each containing a unique m^6^A site in a segment outside of the stem: pri-miR-191, pri-miR-28, and pri-miR-103a.

To ensure that we only observe specific m^6^A modification activity, we tested further the effect of distinct methyltransferase constructs for the representative pri-miRs (Figure 2A). For all three pri-miRs, the highest methylation activity is observed when the methyltransferase domains and the CCCH-type zinc fingers are present (Figure 2B, C). None of the pri-miRs are methylated by the full-length methyltransferase complex with a point mutation of the catalytic residue (D395A). These experiments provide us with multiple METTL3-METTL14 constructs to test their effects on microRNA processing, decoupling the methylation from binding effects.

**Figure 2.**
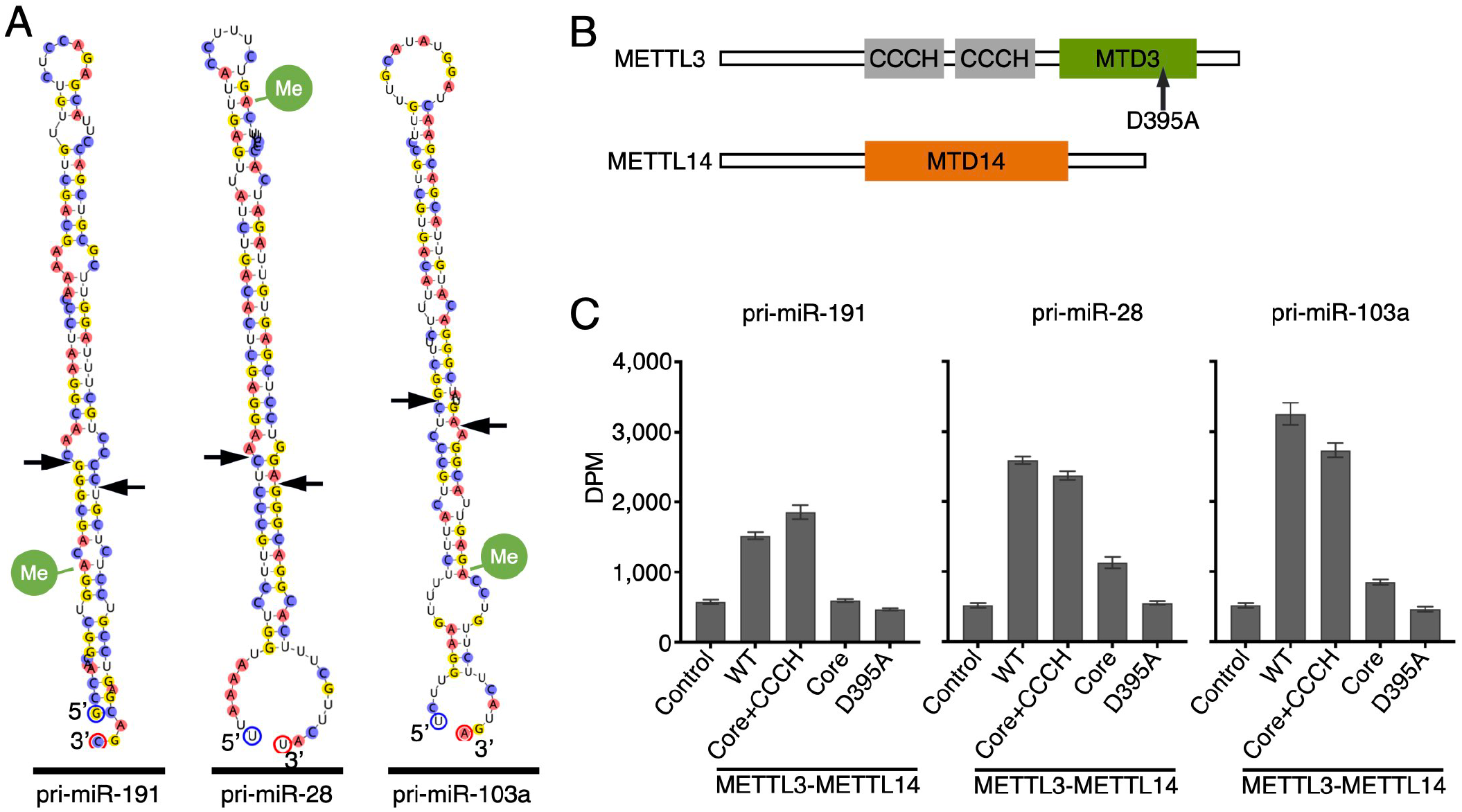
Methylation of pri-miRs by wild-type and mutant METTL3-METTL 14. **(A)** Secondary structure diagrams for pri-miR-191, pri-miR-28 and pri-miR-103a with Microprocessor cut sites shown as arrows. The consensus (DRACH) motifs are indicated with green circles at the expected methylation site. **(B)** Domain organization of METTL3 and METTL 14. **(C)**, *In vitro* methylation assay results for the pri-miRs from (A), for each METTL3-METTL 14 construct. Core contains a complex of MTD3 and MTD14. Core-CCCH contains CCCH domains fused to MTD3 and MTD14.

To test how METTL3-METTL14 affects pri-miR processing by Drosha-DGCR8, we performed staged methylation and processing reactions for each pri-miR (Figure 3A-C). We first added each methyltransferase construct to the pri-miRs with S-adenosylmethionine (SAM) to allow methylation to occur. After pre-incubation with METTL3-METTL14, Microprocessor was then added to the reactions for endonucleolysis to proceed. For all three pri-miRs, the presence of METTL3-METTL14 did not impact the processing efficiency positively or negatively. Furthermore, whether the METTL3-METTL14 constructs are catalytically active or inactive does not affect the impact on the processing efficiency.

**Figure 3.**
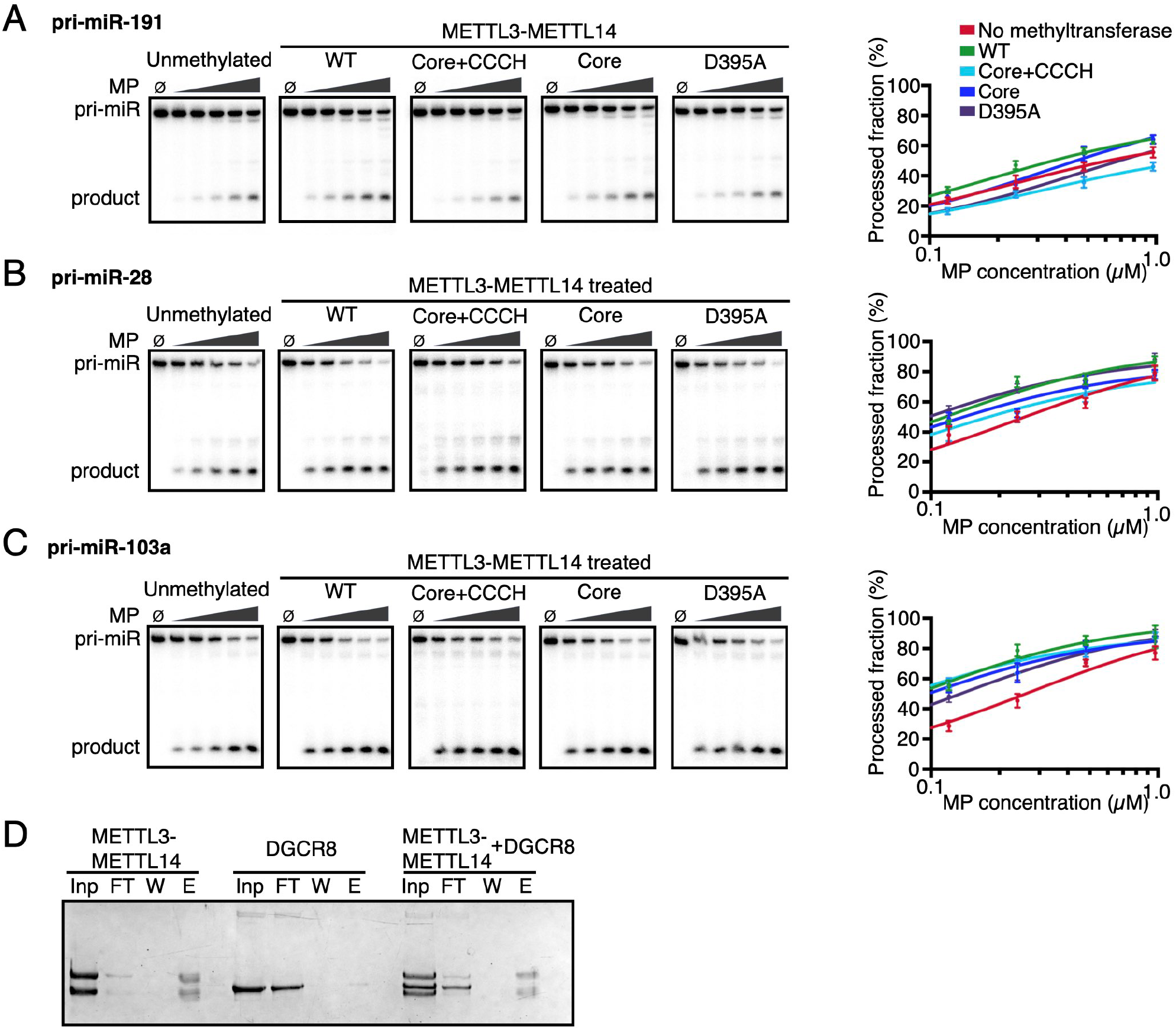
In vitro processing of pri-miRs by Microprocessor in the presence of METTL3-METTL 14. **(A)-(C)** Denaturing PAGE of 5’-radiolabeled pri-miR processing reactions shows different extents of processing with varying Microprocessor levels. For pri-miR-191 **(A)**, pri-miR-28 **(B)**, and pri-miR-103a **(C)**, similar Microprocessor (MP) titrations (2-fold dilutions, 4 to 64 nM; 0, no MP) were performed after pre-incubating the RNAs with 260 nM of the indicated METTL3-METTL 14 construct and 66 μM SAM. Processed fractions were quantified by densitometry and shown as graphs on right. The error bars indicate SD from 3 replicate experiments. **(D)** METTL3-METTL 14 does not form stable complexes with DGCR8 in protein pull-down assays using nickel affinity. SOS-PAGE gel shows input (lnp), flow-through (FT), wash (W) and elution (E) lanes.

The inability of the METTL3-METTL14 constructs to affect Microprocessor activity prompted us to test the leading model where the METTL3-METTL14 binds DGCR8 to recruit Microprocessor to pri-miRs (15,22). We performed a pull-down assay to test if METTL3-METTL14 can form stable complexes with DGCR8 using recombinant proteins (Figure 3D). DGCR8 does not bind METTL3-METTL14 immobilized on beads, suggesting that DGCR8 and METTL3-METTL14 do not make direct contacts sufficient to form stable complexes. Therefore, purified METTL3-METTL14 does not bind DGCR8 and also does not change the efficiency of Microprocessor activity for multiple pri-miRs.

We developed another method to test the effect of the m^6^A modification itself on pri-miR processing. Producing specifically modified long RNAs is challenging, and previous studies have relied on incorporating m^6^A at every occurrence of an adenine in the sequence (14). However, natural modification sites occur more rarely, and substituting every A with m^6^A would dramatically change the RNA structure and chemistry. We established a method to purify large amounts of methylated RNA by using the m^6^A reader protein, YTHDC1 (Figure 4A). We first purified in vitro transcribed pri-miRs. We then treated the pri-miRs with saturating concentrations of METTL3-METTL14 and SAM to maximize methylation extent. The methylated pri-miRs were then allowed to bind the YTH domain of YTHDC1 which binds only the methylated RNAs with high affinity (Figure 4B). Unmethylated RNAs could be washed away when His-tagged YTHDC1 is pulled down with Ni-NTA, and the methylated RNAs could then be extracted away from proteins. Using this enrichment strategy, we were able to produce largely methylated (“m^6^A-enriched”) pri-miRs for miR-28 and miR-103a. Pri-miR-191 was the most difficult to methylate among the three, and our method did not yield enough methylated species. Nevertheless, we successfully enriched for methylated pri-miR-28 and pri-miR-103a, as indicated by YTHDC1-bound bands in gel mobility shift assays (Figure 4C).

**Figure 4.**
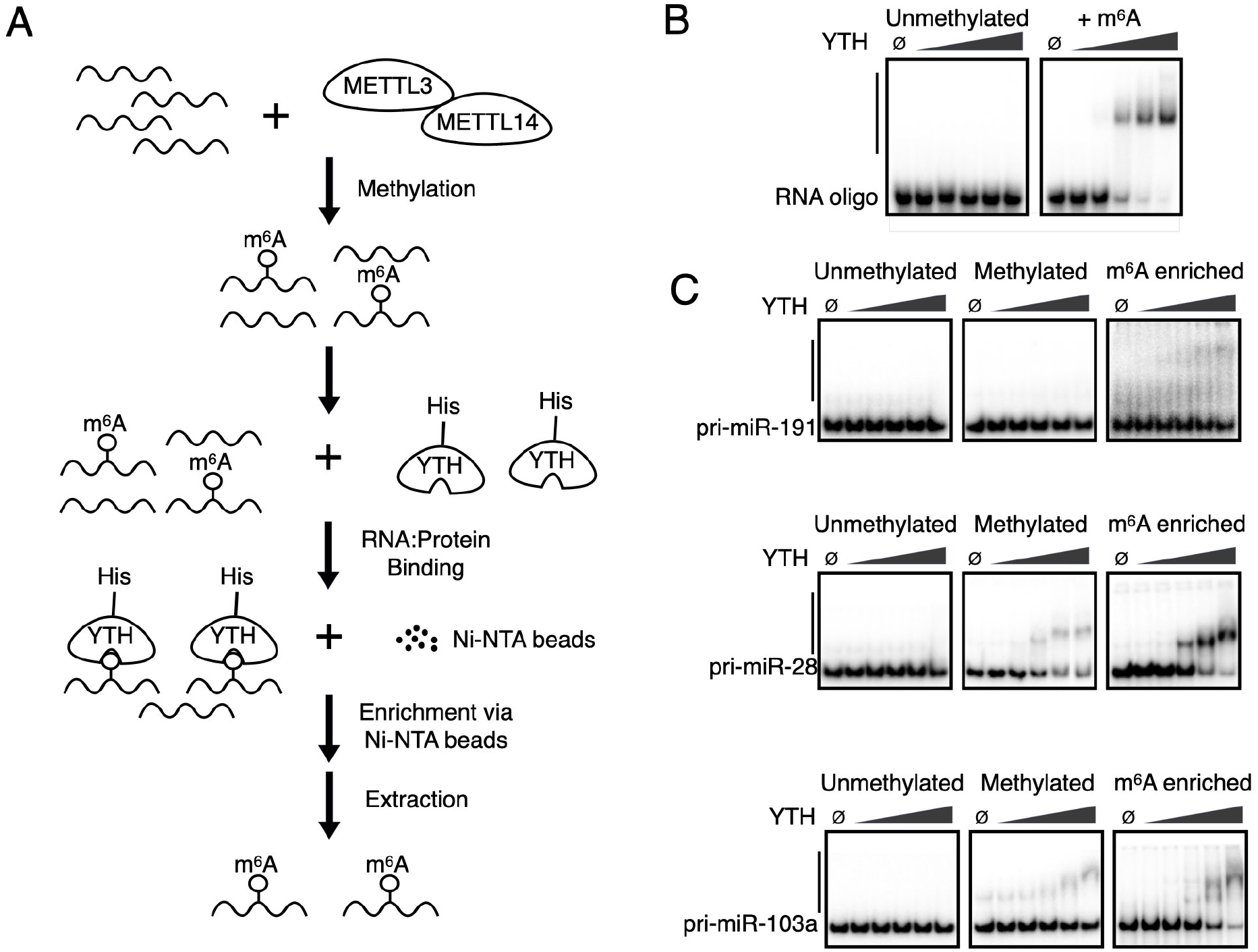
Purification of m^6^A-enriched pri-miRs. **(A)** Strategy to enrich for m^6^A-modified pri-miRs by methylating in vitro transcribed pri-miRs with METTL3-METTL 14 followed by YTHDC1 (YTH)-affinity chromatography and RNA extraction. **(B)** Electrophoretic mobility shift assay (EMSA) shows m6A-specific high affinity of YTH. Synthesized RNA oligonucleotides with the same sequence were used, except for the GGACU motif being replaced with GGm ^**6**^A CU (+ m^**6**^A panel). YTH protein concentrations range from 0.13 μM to 2 μM (2-fold serial dilution; 0, no protein). Solid vertical lines indicate YTH-bound fractions. **(C)** EMSA for m^**6**^A-enriched pri-miRs and YTH shows successful purification of m6A-modified pri-miR-28 and pri-miR-103a. The conditions were identical to **(B)**.

The m^6^A-enriched pri-miRs were compared to the unmethylated counterparts for their ability to be cleaved by Microprocessor (Figure 5A). Titration of the Drosha-DGCR8 complex shows that similar processing activity was observed at each Microprocessor concentration between the two RNA species, suggesting that incorporating m^6^A is not sufficient to impact processing efficiency at variable levels of the processing enzyme.

**Figure 5.**
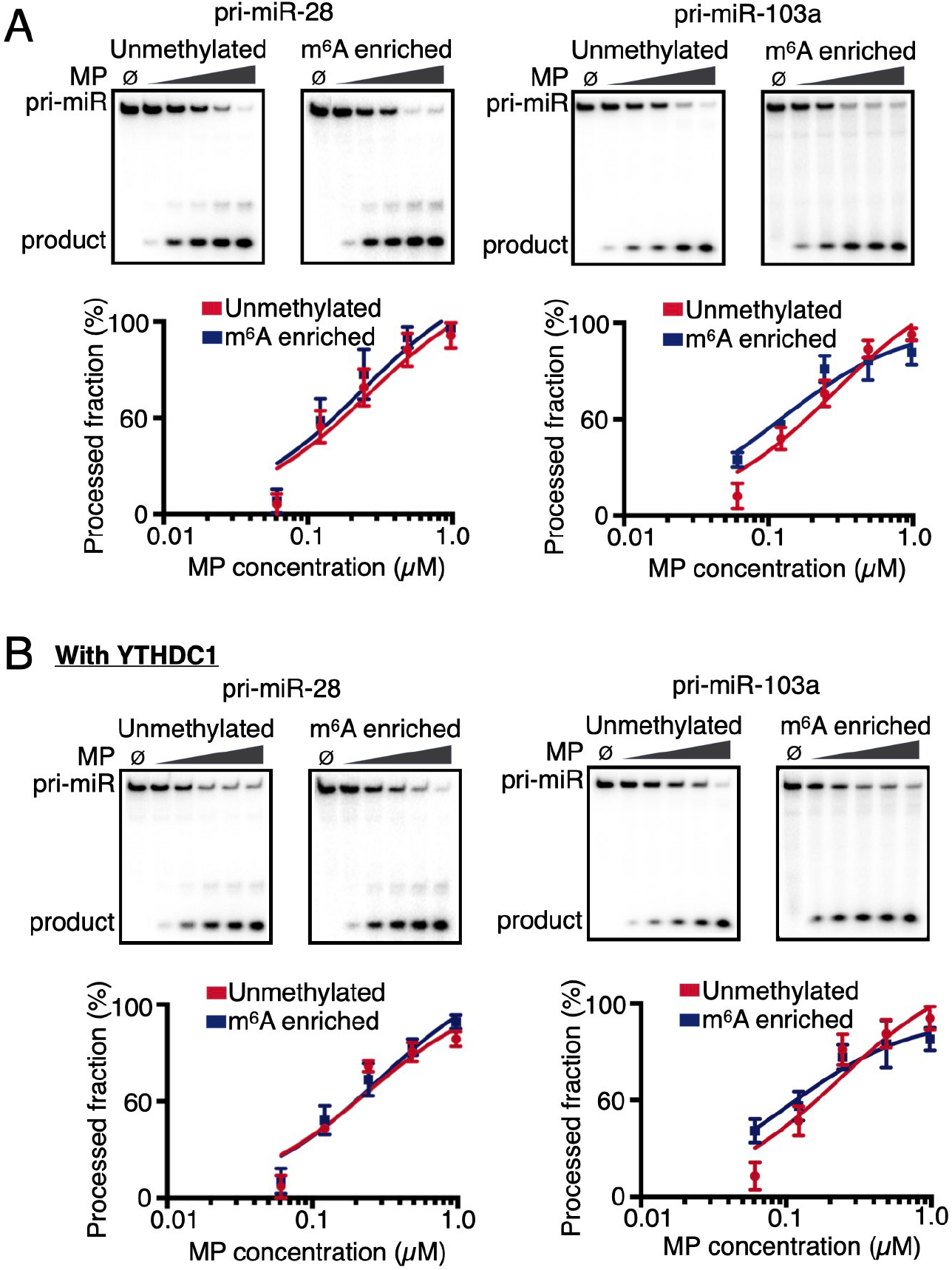
Processing of m^6^A-enriched pri-miRs by Microprocessor. **(A)** In vitro pri-miR processing assays show that methylated and m^6^A-enriched pri-miR-28 and pri-miR-103a are processed similarly to the unmethylated pri-miRs. Microprocessor (MP) titrations (2-fold dilutions over 4 to 65 nM; 0, no MP) were performed. Processed fractions were quantified by densitometry. The error bars indicate SD from 3 replicate experiments. **(B)** Similar to **(A)**, except for the addition of 1 μM purified YTHDC1 in the processing reaction.

Pri-miR processing by Drosha-DGCR8 occurs in the nucleus, and YTHDC1 is a major reader protein in the nucleus that exhibits specific and tight affinity for m^6^A-modified RNAs. We thus tested whether the presence of YTHDC1 changes processing by Microprocessor. In the presence of saturating levels of YTHDC1, processing of pri-miRs was similar for both unmethylated and m^6^A-enriched pri-miRs (Figure 5B). Therefore, even in the presence of a reader protein, pri-miR processing is not altered due to m^6^A modification.

Varying enzyme concentrations may not be enough to capture the differences in processing efficiency between the unmethylated and methylated pri-miRs. Thus, we also performed time course experiments for the processing assays to compare the differences between unmethylated and m^6^A-enriched methylated pri-miRs (Figure 6A). The unmethylated and m^6^A-enriched RNAs have similar processing kinetics, even under single-turnover conditions with excess Microprocessor at saturating levels (Figure 6B). Therefore, m^6^A modification in and of itself may not be sufficient to change processing by Microprocessor in most cases.

**Figure 6.**
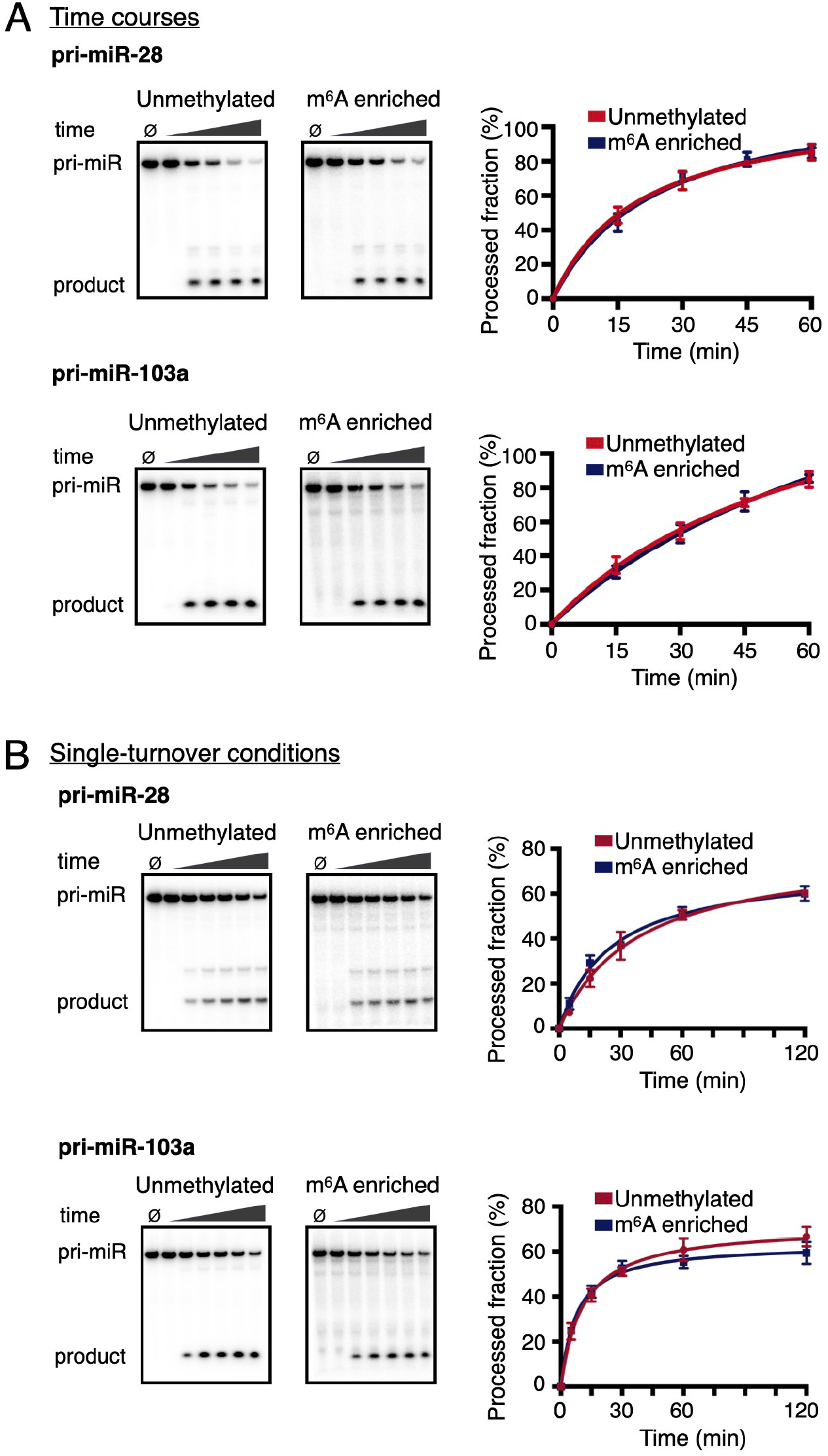
Time-courses of in vitro processing reactions for m^6^A-enriched methylated pri-miRs. **(A)**. In vitro processing of unmethylated and m^**6**^A-enriched pri-miRs (2 nM) by Microprocessor (16 nM) was compared over 0, 15, 30, 45, and 60 minutes (0, no protein control). Processed fractions were quantified by densitometry. The error bars indicate SD from 3 replicate experiments. **(B)** Similar to **(A)**, except for the MP concentration being increased to 260 nM to establish single-turnover conditions.

## Discussion

The question of how specific RNA methyltransferases affect a critical step in microRNA biogenesis is important to determine the crosstalk between RNA modification and processing. Numerous genetic perturbation studies have shown a relationship between microRNA maturation and RNA modification. We show that METTL3 and METTL14 proteins do not directly impact Drosha and DGCR8 activity. The methyltransferase and the processing enzyme complexes do not bind to each other in the absence of other factors. Furthermore, processing efficiencies of pri-miRs are similar in the presence or absence of active or inactive METTL3-METTL14 complexes. Even when we generated mostly methylated pri-miRs via m^6^A-affinity purification, the m^6^A modification was not sufficient to impact microRNA processing by Drosha-DGCR8. Although we cannot exclude the possibility that there may be rare pri-miRs that can be affected by m^6^A, we conclude that modifying pri-miRs with m^6^A is not a general mechanism to broadly promote biogenesis of most microRNAs as observed previously. Therefore, METTL3-METTL14 or m^6^A does not directly enhance microRNA maturation in the absence of other factors or the cellular context.

Microprocessor finds pri-miRs primarily by shape recognition rather than reading base sequences, making secondary structure a key element in substrate detection. Modifying an adenosine with m^6^A in an RNA duplex destabilizes the base-pairs (31), potentially altering the pri-miR structure. However, because METTL3-METTL14 does not readily modify RNA duplexes (Figure 1E), the stem region in the hairpin is not as likely to be modified with m^6^A. Modifications that are more than 20 nucleotides away from the Drosha cut sites may also impact processing. However, given the little effect the sites have within the Microprocessor footprint, the likelihood that pri-miRs are affected directly by the presence of METTL3-METTL14 or m^6^A farther away from the processing machinery is low.

Given that METTL3-METTL14 complex does not seem to directly affect Drosha-DGCR8, the global impact on microRNA levels upon perturbation of the methyltransferase is remarkable. Our studies suggest that the relationship is more complex and may be dependent on other factors. Many studies have suggested that co-transcriptional processing of pri-miRs can influence maturation kinetics. Association of Microprocessor with transcription, splicing factors, and chromatin modifiers tends to promote more efficient processing of the transcripts (32-35). Our studies show that the positive correlation between m^6^A modifications and microRNA processing likely involves such cellular contexts.

## Materials and Methods

### In-vitro transcription and pri-miR purification

Each pri-miR was cloned into a pRZ vector as previously described (36). In vitro transcription and hammerhead and hepatitis delta virus ribozyme cleavage steps were carried out as described in the same reference. The RNAs were purified by denaturing polyacrylamide gel electrophoresis (37).

### Recombinant protein expression and purification

METTL3-METTL14 and METTL1-WDR4 protein complexes were subcloned into pETDuet vectors for coexpression in Rosetta (DE3) pLysS cells (Novagen) in ZYM-5052 media (38). The truncation constructs of METTL3-METTL14 were previously described in (26). Purification was performed as described in previous papers (26,27). Active Microprocessor complex was subcloned into the pFastBacDual vector and Bac-toBac (Thermo) system was used to generate baculoviruses to express the proteins in HighFive (Hi5) cells. Purification was performed as described in (30). The YTH domain of YTHDC1 (residues 345-509) was subcloned into pET21a and expressed in *E. coli* similarly to the methyltransferase complexes. Purification was performed according to the protocol described in (39).

### *In vitro* methylation assay

For in vitro methylation reactions for METTL3-METTL14, each reaction contained 200 nM of RNA, 800 nM of the METTL3-METTL14 complex and 10 μM S-[methyl-^3^H]-adenosyl-L-methionine (SAM) in buffer containing 20 mM Tris pH 7.5, 0.01% Triton-X100, 1 mM DTT, 50 μM ZnCl_2_, 0.2 U/mL RNasin, and 1% glycerol. For the reactions with METTL1-WDR4 protein, each reaction contained 200 nM of RNA, 200 nM of METTL1-WDR4, and 10 μM SAM in the same buffer. Reactions were incubated at 37**°**C for 1 hr, blotted onto Biodyne B nylon membranes, and counted on a scintillation counter (LSC-8000, HITACHI). The incorporation of tritium was measured as disintegrations per minute (DPM).

### Pull-down assay

20 μg of METTL3-METTL14 protein and 20 μg of DGCR8 protein were incubated at 4**°**C for 30 mins in binding buffer (20 mM Tris pH 8, 500 mM NaCl, 20 mM imidazole, 1 mM DTT, 5% (v/v) glycerol), and then applied to Ni-NTA beads. The beads were washed 5 times in the same buffer and eluted with the same buffer with increased (400mM) imidazole.

### Electrophoretic mobility shift assay (EMSA)

RNA oligonucleotides derived from MALAT1 (5’-AACUU AAUGU UUUUG CAUUG GACUU UGAGUU -3’) were synthesized with or without m^6^A modification (Sigma). Oligonucleotides and the indicated pri-miRs were radiolabeled at 5’ ends with γ^32^P-ATP using T4 polynucleotide kinase. Serially diluted YTHDC1 protein (2-fold, 0.13 μM – 2 μM) was incubated with 2 nM RNA in a binding buffer (20 mM Tris pH 8.0, 150 mM NaCl, 10 mM DTT, 50 μM ZnCl_2_, 5% (v/v) glycerol, 100 ng/μL yeast tRNA). The mixtures were analyzed by native polyacrylamide gel electrophoresis before detection using a phosphorimager (Typhoon FLA 9500).

### In vitro pri-miR processing assay

The pri-miRs were radiolabeled at 5’ ends with ^32^P-ATP using T4 polynucleotide kinase. For Microprocessor titrations, the Microprocessor complex (0.004 – 0.0651 μM, 2-fold serial dilutions) was added to 2 nM RNA in a reaction buffer containing 30 mM Tris pH 7.5, 5 mM DTT, 10 mM MgCl_2_, 3% glycerol, and 0.2 U/mL RNasin, and the reaction was incubated at RT for 30 min. For pre-incubation with METTL3-METTL14, each reaction was first treated with 260 nM METTL3-METTL14 and 66 μM SAM in the same buffer before adding Microprocessor. When indicated, 1.04 μM YTHDC1 was added and allowed to bind for 30 min before adding Microprocessor. For the first time-course experiment, each reaction contained 2 nM RNA, 10 μM SAM, and 16 nM of Microprocessor in the same reaction buffer. For the single-turnover time course, the concentration of Microprocessor was increased to 260 nM and the reaction was carried out at 12 °C. All reactions were quenched with 1% SDS, 70mM EDTA, and proteinase K before denaturing polyacrylamide gel electrophoresis and detection by a phosphorimager.

### Methylation and enrichment of pri-miRNA

Each pri-miR (200 nM) was incubated with 800 nM METTL3-METTL14 at 37**°**C for 1 hr in a buffer solution containing 20 mM Tris pH 7.5, 0.01% Triton X-100, 50 μM ZnCl_2_, 1 mM DTT, 1% (v/v) glycerol, 0.2 U/mL RNasin, and 10 μM SAM. For enrichment, 150 nM of pre-methylated pri-miR was incubated with 25 μM of YTHDC1 protein for 2 hrs at 4**°**C in a binding buffer (20 mM Tris pH 8, 150 mM NaCl, 1 mM DTT). After incubation, the RNA-protein mixture was added to Ni-NTA beads and incubated for 2 hrs at 4**°**C. The mixture was then washed with buffer and eluted with the same buffer containing 250mM imidazole. Each m^6^A-enriched pri-miR batch was verified by EMSA with YTHDC1 as described above.

## Data Availability

All data presented in the work are available within the main figures and the supplementary data.

## Supplementary Data

Supplementary Table S1

## Acknowledgments

The work was supported by NIH/NIGMS (R01GM122960 to YN), NIH/NCI (R01CA258589 to YN), and Welch Foundation (I-2115-20220331 to YN). Y.N. is a Southwestern Medical Foundation Scholar in Biomedical Research, UT Southwestern Presidential Scholar, Doris and Bryan Wildenthal Distinguished Chair in Medical Science, Pew Biomedical Scholar (27339), and Packard Fellow (2013-39275).

---

**Supplementary Table S1.** List of pri-miRs featuring consensus (DRACH) motifs within 20 nucleotides upstream and 20 nucleotides downstream of Drosha cut sites. These pri-miRs contain mature microRNAs that were significantly suppressed upon METTL3 knock-down (fold change <=0.5 and p <=0.05) (Data adapted from Alarcon, et al, 2015, Nature).

|  | Name | Fold change kD/Control | P value | DRACH motif within 20/20 |
| --- | --- | --- | --- | --- |
| 1 | hsa-miR-146a-5p | 0.30087759 | 0.00248 | 2 |
| 2 | hsa-miR-193b-3p | 0.149060833 | 0.00552 | 2 |
| 3 | hsa-miR-130a-3p | 0.191415937 | 0.00563 | 0 |
| 4 | hsa-miR-92b-3p | 0.265886434 | 0.00604 | 2 |
| 5 | hsa-miR-138-5p | 0.161751101 | 0.00687 | 0 |
| 6 | hsa-miR-29b-1-5p | 0.38926631 | 0.00705 | 1 |
| 7 | hsa-miR-485-3p | 0.495153177 | 0.00767 | 0 |
| 8 | hsa-miR-660-5p | 0.1978095 | 0.00856 | 2 |
| 9 | hsa-let-7d-3p | 0.31436026 | 0.00905 | 1 |
| 10 | hsa-miR-93-5p | 0.173131382 | 0.00941 | 2 |
| 11 | hsa-miR-22-3p | 0.294217484 | 0.01262 | 2 |
| 12 | hsa-miR-342-3p | 0.165957901 | 0.01357 | 2 |
| 13 | hsa-miR-103a-3p | 0.159456462 | 0.01464 | 1 |
| 14 | hsa-miR-130b-3p | 0.157730623 | 0.01691 | 0 |
| 15 | hsa-miR-106b-5p | 0.166780598 | 0.0171 | 1 |
| 16 | hsa-miR-30a-3p | 0.313994591 | 0.01929 | 3 |
| 17 | hsa-miR-24-3p | 0.310754475 | 0.01945 | 2 |
| 18 | hsa-miR-30d-5p | 0.311085265 | 0.01953 | 3 |
| 19 | hsa-miR-425-5p | 0.147363945 | 0.02 | 1 |
| 20 | hsa-miR-331-3p | 0.118433361 | 0.0205 | 2 |
| 21 | hsa-miR-365a-3p | 0.158419498 | 0.02077 | 1 |
| 22 | hsa-miR-19b-3p | 0.155877283 | 0.02141 | 3 |
| 23 | hsa-miR-30a-5p | 0.368490855 | 0.02144 | 3 |
| 24 | hsa-miR-221-3p | 0.49863549 | 0.02149 | 2 |
| 25 | hsa-miR-126-3p | 0.131641794 | 0.02225 | 3 |
| 26 | hsa-miR-99b-5p | 0.190733083 | 0.02321 | 2 |
| 27 | hsa-miR-191-5p | 0.284144088 | 0.0245 | 1 |
| 28 | hsa-let-7a-2-3p | 0.208313385 | 0.02937 | 3 |
| 29 | hsa-miR-15b-5p | 0.318185046 | 0.03084 | 0 |
| 30 | hsa-miR-16-2-3p | 0.37904403 | 0.0318 | 3 |
| 31 | hsa-miR-92a-3p | 0.388160942 | 0.03205 | 2 |
| 32 | hsa-miR-193a-5p | 0.225199689 | 0.03212 | 1 |
| 33 | hsa-miR-30c-5p | 0.429266126 | 0.03259 | 3 |
| 34 | hsa-miR-455-3p | 0.126870246 | 0.03654 | 1 |
| 35 | hsa-miR-192-5p | 0.361675904 | 0.03814 | 3 |
| 36 | hsa-miR-138-1-3p | 0.387681942 | 0.03874 | 0 |
| 37 | hsa-miR-10b-5p | 0.234081502 | 0.03948 | 2 |
| 38 | hsa-miR-128-3p | 0.152475037 | 0.04171 | 1 |
| 39 | hsa-miR-28-5p | 0.25599257 | 0.04188 | 1 |
| 40 | hsa-miR-30e-5p | 0.418652297 | 0.0419 | 2 |
| 41 | hsa-miR-106a-5p | 0.257606899 | 0.04248 | 0 |
| 42 | hsa-miR-140-3p | 0.263818853 | 0.04261 | 2 |
| 43 | hsa-miR-7-5p | 0.225879857 | 0.04296 | 2 |
| 44 | hsa-miR-15a-5p | 0.243099271 | 0.04425 | 1 |
| 45 | hsa-miR-20a-5p | 0.274239383 | 0.04441 | 2 |
| 46 | hsa-miR-183-5p | 0.173437481 | 0.04461 | 4 |
| 47 | hsa-miR-210-3p | 0.220451925 | 0.04476 | 2 |
| 48 | hsa-miR-17-5p | 0.248733 | 0.04619 | 2 |
| 49 | hsa-miR-25-3p | 0.366817595 | 0.04655 | 4 |
| 50 | hsa-miR-320a | 0.267294265 | 0.04921 | 1 |

